# An open-source, ready-to-use and validated ripple detector plugin for the Open Ephys GUI

**DOI:** 10.1101/2022.04.01.486754

**Authors:** Bruno Monteiro de Sousa, Eliezyer Fermino de Oliveira, Ikaro Jesus da Silva Beraldo, Rafaela Schuttenberg Polanczyk, João Pereira Leite, Cleiton Lopes Aguiar

## Abstract

Sharp wave-ripples (SWRs, 100-250 Hz) are oscillatory events extracellularly recorded in the CA1 subfield of the hippocampus during sleep and quiet wakefulness. SWRs are thought to be involved in the dialogue between the hippocampus and cortical regions to promote memory consolidation during sleep and memory-guided decision making. Many studies employed closed-loop strategies to either detect and abolish SWRs within the hippocampus or manipulate other relevant areas upon ripple detection. However, the code and schematics necessary to replicate the detection system are not always available, which hinders the reproducibility of experiments among different research groups. Furthermore, information about performance is not usually reported. Here, we present the development and validation of an open-source, real-time ripple detection plugin integrated into the Open Ephys GUI. It contains a built-in movement detector based on accelerometer or electromyogram data that prevents false ripple events (due to chewing, grooming, or moving, for instance) from triggering the stimulation/manipulation device. To determine the accuracy of the detection algorithm, we first carried out simulations in Matlab with synthetic and real ripple recordings. Using a specific combination of detection parameters (amplitude threshold of 5 standard deviations above the mean, time threshold of 10 ms, and RMS block size of 7 samples), we obtained a 97% true positive rate and 2.48 false positives per minute on the real data. Next, an Open Ephys plugin based on the same detection algorithm was developed, and a closed-loop system was set up to evaluate the round trip (ripple onset-to-stimulation) latency over synthetic data. The lowest latency obtained was 34.5 ± 0.5 ms. Besides contributing to increased reproducibility, we anticipate that the developed ripple detector plugin will be helpful for many closed-loop applications in the field of systems neuroscience.

## 1. INTRODUCTION

Sharp wave-ripples – electrophysiological oscillations in the local field potential (LFP) of the hippocampal CA1 region – are thought to coordinate the repetitive reactivation of memory-related ensembles and contribute to the integration and stabilization of memory representations (Buzsáki, 2015). Regarding their oscillatory features, most SWR events last from 50 to 150 ms, and their frequency components are within the 100-250 Hz range (Buzsáki, 2015; Dutta, Ackermann and Kemere, 2019; Fernández-Ruiz et al., 2019; Adamantidis, Herrera and Gent, 2019). Studies that investigate the causal link between SWRs, learning, and memory consolidation during sleep have usually employed closed-loop, real-time ripple disruption experiments using electrical stimulation (Girardeau et al., 2009; Ego-Stengel and Wilson, 2010; Maingret et al., 2016) or, more recently, optogenetic techniques (Kovács et al., 2016; Norimoto et al., 2018; Oliva et al., 2020). Closed-loop strategies have also been used to disrupt SWRs during quiet wakefulness, a period in which the awake animal presents slow movement or immobility (Jadhav et al., 2012; Roux et al., 2017). The module responsible for the real-time ripple detection is often contained within proprietary, dedicated hardware (Table I). Additionally, the detailed schematics and source code necessary to build a similar detection setup are rarely readily available. This makes it difficult to reproduce results obtained across different laboratories.

**Table I.** Overview of experimental details in studies using closed-loop systems based on ripple detection. For other related information, see Aleman-Zapata, van der Meij and Genzel, 2021.

| Authors | Animal model | SWR recording region | Online ripple detection components | Detection codes/circuits are publicly available? | Graphical interface for adjusting the detection parameters? | Ripple-triggered manipulation | Detection accuracy and round trip latency | Manipulation disabled during wakefulness/movement? |
| --- | --- | --- | --- | --- | --- | --- | --- | --- |
| Girardeau et al. (2009) | Rats | CA1 pyramidal layer (dorsal hippocampus) | Power1401 acquisition system + Spike2 software (CED) | No | Not found | Single-pulse commissural electrical stimulation | Not found | Not found |
| Ego-Stengel and Wilson (2010) | Rats | CA1 pyramidal layer | KrohnHite3384 analog filters and FHC Window Discriminator (threshold crossing) | Doesn't apply | Doesn't apply | Burst of biphasic electrical pulses on CA1 afferents | Not found | Not found |
| Jadhav et al. (2012) | Rats | CA1 (dorsal hippocampus) | NSpike data acquisition system | No | Not found | Biphasic, single-pulse commissural electrical stimulation | Ripple oscillation terminated within 25 ms of SWR onset | Yes - online speed filter prevented false positive detections during high speed periods (threshold from 5 to 10 cm/s) |
| Nokia et al. (2012) | Rabbits | Right hippocampus | Custom-made hardware | Yes | Doesn't apply | Light pulses from three LEDs placed in front of the rabbit | 89% of SWR events identified offline were detected online | No |
| Stark et al. (2014) | Mice/Rats | CA1 pyramidal layer | Programmable Digital Signal Processor RX6 + RPvdsEx (Tucker-Davis Technologies) | No | Not found | Optogenetic manipulations | Median detection lag: 15 ms; range 12–24 ms; median false negative rate, 27%; range: 7%–45%; Bayes error of the classifier: 16% (median; range: 9%–28%) | Not found |
| de Lavilléon et al. (2015) | Mice | CA1 pyramidal layer | Power1401 acquisition system + Spike2 software (CED) | No | Not found | Train of electrical pulses | The authors reported spike sorting performance | Not found |
| Papale et al. (2016) | Rats | CA1 layer (dorsal hippocampus) | NSpike data acquisition system (Jadhav et al., 2012) | No | Not found | Biphasic, single-pulse electrical stimulation | Not found | Not found |
| Novitskaya et al. (2016) | Rats | CA1 pyramidal layer (dorsal hippocampus) | Power1401 acquisition system + Spike2 software (CED) | No | Not found | Trains of biphasic electrical pulses | Not found | Not found |
| Maingret et al. (2016) | Rats | CA1 pyramidal layer | Power1401 acquisition system + Spike2 software (CED) | No | Not found | Single-pulse electrical stimulation | Not found | Not found |
| Kovács et al. (2016) | Mice | CA1 pyramidal layer (dorsal hippocampus) | Custom-made hardware (Nokia et al. 2012) | Yes | Doesn't apply | Optogenetic manipulations | Not found | No |
| Talakoub et al. (2016) | Rhesus monkeys (Macaca mulatta) | CA1/2 and CA3/DG | Programmable Digital Signal Processor HPP (Neuralynx, Inc) + FPGA | No | Not found | Biphasic, single-pulse electrical stimulation | FPR estimate: 12% | No - but artifacts in control electrodes outside the hippocampus blocked ripple detection |
| van de Ven et al. (2016) | Mice | CA1 pyramidal layer (dorsal hippocampus) | Dual-stage amplifier (Sensorium Inc.) + computer with real-time kernel | No | Not found | Optogenetic manipulation | True positive: 80.1% ± 1.0%; Mean Latency: 7.68 ± 0.30 ms before ripple peak power | No - but artifacts in control electrodes outside the hippocampus blocked ripple detection |
| Roux et al. (2017) | Mice | CA1 pyramidal layer (dorsal hippocampus) | Programmable Digital Signal Processor RX6 + RPvdsEx (Tucker-Davis Technologies) (Stark et al. 2014) | No | Not found | Optogenetic manipulation | TPR: 83 ± 4%; FPR: 63 ± 4%; Detection latency: median ± SD = 16 ± 20 ms | No |
| Rangel Guerrero et al. (2018) | Mice | CA1 pyramidal layer (dorsal hippocampus) | Custom-made hardware (Nokia et al. 2012) | Yes | Doesn't apply | Optogenetic manipulation | Not found | No |
| Dutta, Ackermann and Kemere (2019) | Rats | CA1 | Open Ephys and SpikeGadgets acquisition systems | Partially (analysis codes available; codes for online detection not identified) | Yes | Digital pulse from a BeagleBone microcontroller | Data acq. latency: 7.5-13.8 ms and 1.35-2.6 ms for USB and ethernet, respectively. Algorithmic latency approx. 20-66ms, TPR > 95% and <10 FP/min, depending on the parameters. | No |

**Table I - (Continued)**
| Authors | Animal model | SWR recording region | Online ripple detection components | Detection codes/circuits are publicly available? | Graphical interface for adjusting the detection parameters? | Ripple-triggered manipulation | Detection accuracy and round trip latency | Manipulation disabled during wakefulness/movement? |
| --- | --- | --- | --- | --- | --- | --- | --- | --- |
| Norimoto et al. (2018) | Mice | CA1 pyramidal layer | Programmable Digital Signal Processor RX6 + RPDsEx (Tucker-Davis Technologies) (Stark et al. 2014) | No | Not found | Optogenetic manipulation | 23 ± 12 ms delay for stimulation after ripple onset; FPR: 19.2 ± 1.0%; False negative rate: 2.3 ± 1.8% | No |
| Michon et al. (2019) | Rats | CA1 pyramidal layer | DigiLynx acquisition system + Falcon software | No | Not found | Biphasic electrical stimulation | S3 figure in Michon et al. 2019 | No - but hippocampal ripple detections were ignored if they fell around spurious detections in the cortical signals |
| Fernández-Ruiz et al. (2019) | Rats | CA1 pyramidal and stratum radiatum layers | Programmable processor + Spike2 software (CED) | Partially (analysis codes available; codes for online detection not found) | Not found | Optogenetic manipulation | Not found | Yes - ripples detected only during immobility (speed < 5 cm/s) and low-theta power periods |
| Oliva et al. (2020) | Mice | CA2 | Programmable processor (CED) + custom-made hardware | No | Not found | Optogenetic manipulation | Not found | No - but artifacts in control electrodes outside the hippocampus blocked ripple detection |
| Igata, Ikegaya and Sasaki (2020) | Rats | CA1 | Cerebus recording system + custom created C code | No | Not found | Single-pulse electrical stimulation | Not found | Yes - ripples detected only during immobility (speed < 5 cm/s) |
| Gridchyn et al. (2020) | Rats | CA1 pyramidal layer | Data acquisition system (Axona Ltd) + Python software | Yes | No | Optogenetic manipulation | Reaction time (from first spike to LFP deflection caused by the light pulse): 1.04 ms ± 0.09 ms | Not found |
| Aleman-Zapata, Morris and Genzel. (2020, Preprint) | Rats | CA1 pyramidal layer | Open Ephys acquisition system + Cython | Yes | Yes | Electrical stimulation | Not found | Not found |

It is also worth mentioning that not all studies report the accuracy and latency of ripple detection (Table I). If the reason is the non-execution of prior performance tests, investigators may be inadvertently applying an excess stimulation due to more false-positive events. In some cases, closed-source solutions may limit the feasibility of tests to evaluate the effects produced by variations in the detection parameters, and make it challenging to execute a system-level analysis of the detection performance. Therefore, the trade-offs between performance and over-stimulation side-effects, such as tissue damage, seizures, and plasticity induction, may not be fully understood and controlled by the investigator (Dutta, Ackermann and Kemere, 2019).

Valuable efforts have been applied to provide a real-time ripple detector whose closed-loop, system-level performance is known so that investigators can be aware of the limitations and possibilities offered by such a tool (Dutta, Ackermann and Kemere, 2019). Nevertheless, the scientific community still lacks open-source, publicly available ripple detection modules already integrated with widely used electrophysiological recording platforms. Additionally, the increasingly common long-lasting recordings *in vivo* demand built-in, automatic strategies capable of restricting ripple-triggered experimental interventions only to periods of low movement (including sleep and quiet wakefulness), when ripple events are most likely to occur. Movement artifacts during the active phase of rats and mice, for instance, occur frequently and can result in wrongly-triggered stimulation, interfering with the experimental protocol. Although most of the closed-loop studies on SWRs carry out interventions only during the initial periods of sleep, new evidence shows that associated replay activity occurs over a much longer timespan (Giri et al., 2019). Moreover, even in experimental protocols in which stimulation is applied only during the first few hours of the post-learning sleep, it is expected that the animal presents periods of wakefulness due to the polyphasic nature of its sleep-wake cycle (Mong and Cusmano, 2016).

We developed a ready-to-use, real-time ripple detector plugin for the Open Ephys GUI (Siegle et al., 2017). The built-in algorithm to detect movement based on electromyogram (EMG) or accelerometer data offers the possibility to prevent ripple events from being detected during periods of high activity, where they are likely to be false alarms. A 3-axis accelerometer is already embedded in many headstage models commonly used in conjunction with the Open Ephys acquisition board; thus, movement detection can be performed without using additional implants or a video tracking apparatus. First, to evaluate the dependence of our algorithm’s detection accuracy on multiple combinations of parameters, we conducted an offline Matlab (MathWorks) simulation of the plugin over synthetic ripple data and pre-recorded real hippocampal CA1 data (hc-3 dataset, Mizuseki et al., 2013). Next, to evaluate the round trip (event-to-stimulation) latency, a closed-loop architecture similar to the one used in real-time experiments was arranged. In this case, the living animal (or tissue) is replaced by the synthetic data converted to an audio signal and connected directly to an Intan headstage. This closed-loop structure also made it possible to examine the effects of varying other influencing parameters, i.e., the data buffer length and the number of samples considered in the RMS calculation. To exemplify the mechanism’s functioning that blocks ripple events when movement is detected, we present a segment of real accelerometer data and the corresponding periods of silencing for different combinations of parameters. The accelerometer data was obtained from a publicly available dataset (TingleyD dataset, Petersen, Hernandez and Buzsáki, 2020). Finally, although the developed plugin does not perform multichannel ripple detection, various instances of the module can be combined to build a multichannel logic that results in reduced false-positive rates.

## 2. METHODS

### 2.1. Implementation of the detection algorithm

The ripple detector was implemented in C++ as a plugin for the Open Ephys GUI v0.5.5 (**Fig. 1a**). We used the Microsoft Visual Studio 2019 platform to develop and debug the code. Researchers interested in using the ripple detector must download the source code and compile the plugin following the instructions available in the Developer Guide section at open-ephys.github.io/gui-docs/. The plugin’s source code is publicly available at github.com/bmsousa91/RippleDetector. The synthetic dataset used in sections **3.1** and **3.3**, and more details on how to use the plugin, are available on the same website.

**Figure 1.**
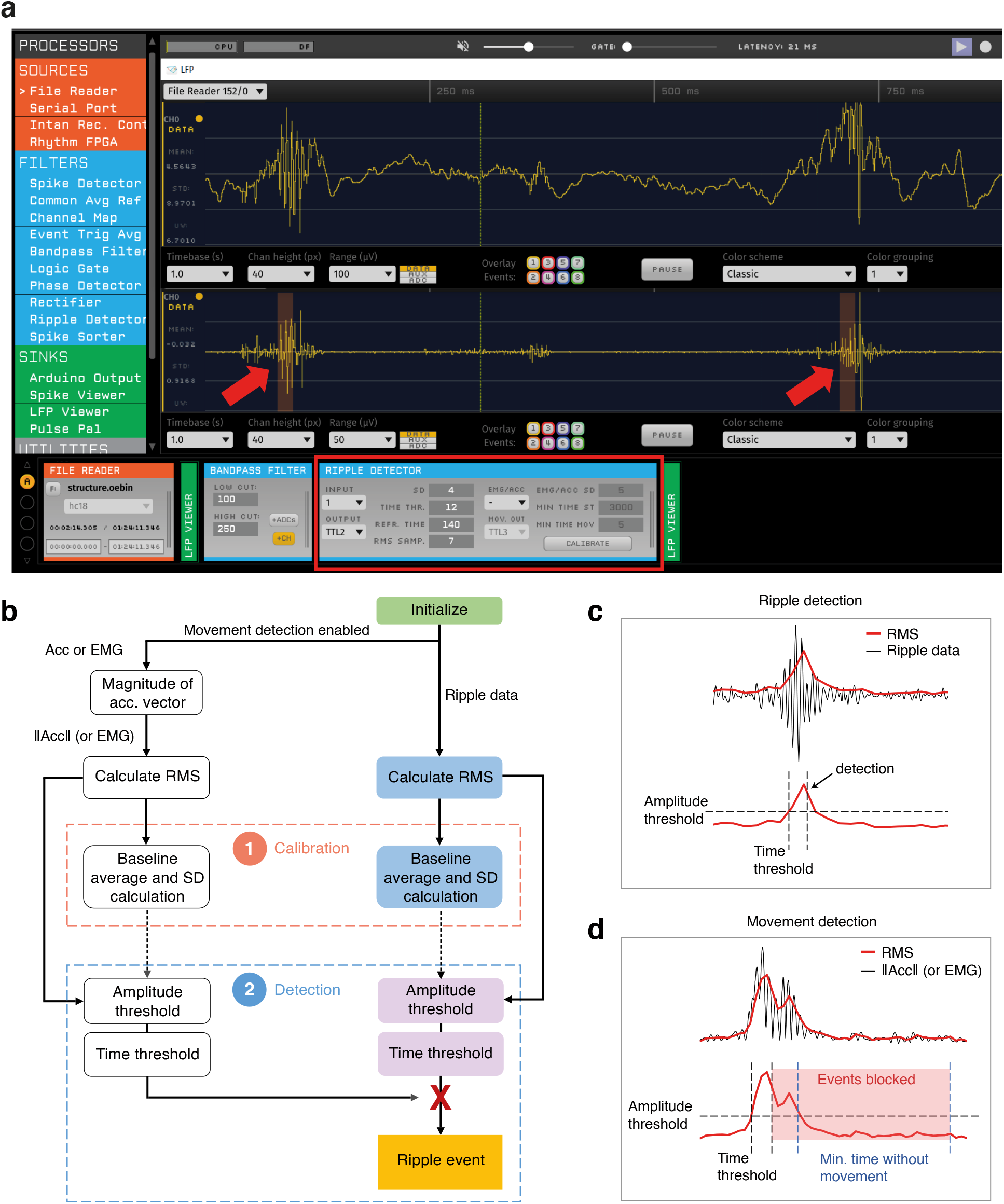
Ripple detector plugin for the Open Ephys platform. **a)** Through the plugin’s graphical interface (highlighted in red), users can adjust the detection parameters. The red arrows indicate detected ripple events. The data shown in this figure were downloaded from the CRCNS website (hc-18 dataset – Drieu, Todorova and Zugaro, 2018a; Drieu, Todorova and Zugaro, 2018b). **b)** Step-by-step flowchart of the detection algorithms. Data must be previously filtered in the ripple band. **c)** Illustrative example of RMS threshold crossing and ripple detection. Continuous black line: filtered ripple data; red line: RMS envelope of the filtered data; dashed lines: amplitude and time thresholds for ripple detection. **d)** Illustrative example of movement detection. Continuous black line: filtered accelerometer or EMG data; red line: RMS envelope; dashed black lines: amplitude and time threshold for movement detection; dashed blue lines: define the minimum period of immobility to restore the plugin’s ability to raise detection events. The area in red represents the period in which ripple events are silenced due to motion detection.

### 2.2. Ripple detection algorithm and movement-driven silencing of events

Real-time detection algorithms must operate with minimal delay while maintaining acceptable accuracy levels. Furthermore, investigators benefit from a more straightforward detection logic since the effects of adjusting specific parameters are more predictable. Aiming at those features, our ripple detection algorithm was implemented based on the root mean square (RMS) of the filtered signal (**Fig. 1b**). The ripple detector does not perform any filtering internally and therefore requires that signals are pre-filtered in the ripple frequency band, which may be achieved by using the built-in Bandpass Filter module (second-order Butterworth filter) in the Open Ephys GUI. The RMS values of the incoming samples provide a rough envelope of the ripple-band filtered data (they can also be thought of as a recurring estimation of the standard deviation if the mean is considered zero). Initially, a calibration period – the first 20 seconds of recording – defines the data baseline, when RMS mean and standard deviation are calculated. Subsequently, RMS values are compared against amplitude and time thresholds during the online detection stage. The amplitude threshold is defined by a fixed number of standard deviations above the mean. RMS values must exceed the amplitude threshold and remain above it for a minimum period – defined as the time threshold – to raise a ripple event (**Fig. 1c**). The time threshold reduces false detections due to brief periods of RMS augmentation in the ripple frequency band that do not correspond to true ripple oscillation (e.g., noise artifacts).

The movement detection algorithm (**Fig. 1b**) is very similar to the ripple detection logic. It operates on filtered EMG or accelerometer data to disable ripple events when movement is detected. If the accelerometer is selected to detect movement, the magnitude of the 3D acceleration vector is calculated before the RMS computation. Ripple events are blocked when the RMS values cross the corresponding amplitude and time thresholds. Reactivating detection requires a minimum period of immobility (i.e., a sequence of RMS values below the amplitude threshold) (**Fig. 1d**).

### 2.3. Synthetic ripple data

Synthetic data was created by adding simulated ripple, fast ripple, and spike events to a pink noise background signal (**Fig. 2a**). The corresponding features to simulate those events were extracted from prior characterization studies (Buzsáki et al., 1992; Bragin et al., 1999; Nguyen et al., 2009; Buzsáki, 2015). Ripples were represented as pure sinusoidal signals whose frequency distribution followed a Gaussian process with μ = 200 Hz and σ = 25 Hz. Its amplitude was limited by a sinusoidal envelope whose duration was determined by a Gaussian process with μ = 100 ms and σ = 25 ms. The signal-to-noise ratio (SNR) was around 18 dB. Fast ripples were thought to simulate higher frequency activity in the hippocampus (e.g., seizure onset). They were generated similarly to the ripples but with a shorter duration (μ = 60 ms; σ = 15 ms), higher amplitude, and increased frequency content (μ = 425 Hz; σ = 50 Hz). Finally, artificial spikes were synthesized as Gaussian curves with high amplitude and low standard deviation (σ = 0.0003 ms) to simulate a brief and expressive increase in the ripple-band power. The non-ripple events (fast ripples and spikes) test the algorithm’s ability to reject spurious activities that contaminate the ripple frequency band correctly. The final duration of the synthetic data was 10 minutes (30 kHz sampling rate), and one of those three events was injected every 1.5 s, with a likelihood of 33.3% each. There were 400 events: 139 ripples, 124 fast ripples, and 137 spikes.

**Figure 2.**
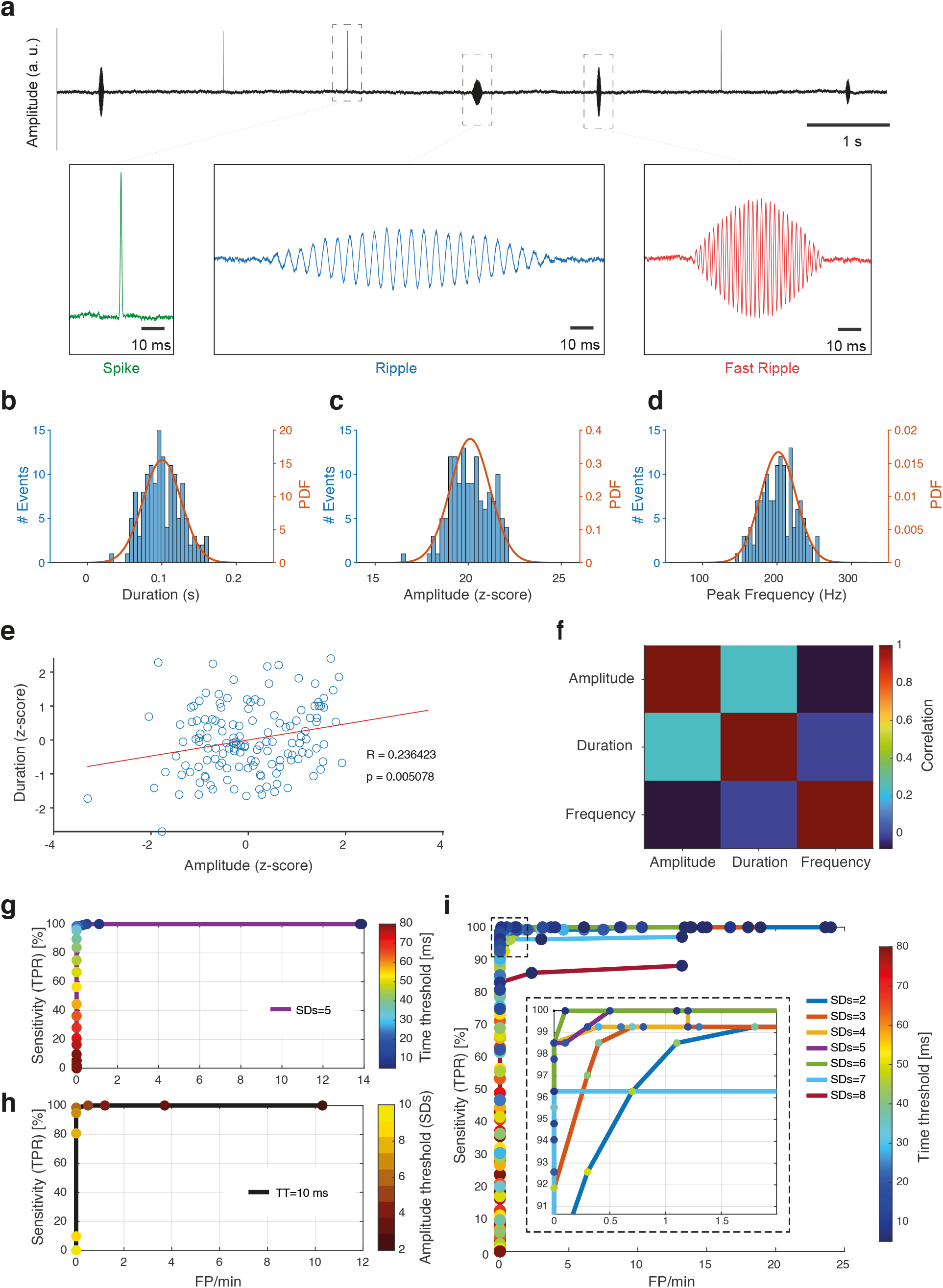
Characterization of synthetic ripple data and corresponding ROC curves for offline simulations of the plugin in Matlab. **a)** Example of a synthetic data segment. Simulated events (spikes, ripples, and fast ripples) are inserted into a pink noise background signal. **b)** Duration of ripple simulated events (n=139 events, Kolmogorov-Smirnov normality test with p=0.5206; mean=100.4 ms, SD=25.6 ms). **c)**

### 2.4. Hippocampal CA1 recordings from the hc-3 dataset

To evaluate the plugin’s performance over real ripple recordings (section **3.2**), pre-recorded hippocampal CA1 data from the hc-3 dataset were downloaded at the CRCNS website (crcns.org/data-sets/hc/hc-3). The hc-3 dataset contains LFP and multiunit recordings from different hippocampal regions of male Long Evans rats (3-8 months old, 250-400 g) while animals performed several tasks (Mizuseki et al., 2013; Mizuseki et al., 2014). The authors used 4-, 8-, 12- or 16-shank silicon probes, each shank containing 8 recording sites (160 µm^2^ each site, impedance of 1-3 MΩ). Recording sites formed a two-dimensional arrangement and were separated by 20 µm vertically. We considered LFP recordings from the animal ec014, day 42, during the sleep sessions ec014.42.794, ec014.42.796, and ec014.42.798 (**Fig. 3a**). Data from each sleep section was concatenated to result in a 30-minutes long single recording (1250 Hz sampling rate). In all experiments using the hc-3 dataset, the selected recordings were captured from the silicon probe shank 1, channel 8 (indicated by the dataset providers as the shank 1 channel with the most pronounced ripples). To test the multichannel ripple detection strategy (section **3.5**), channels 6 and 4 from the same shank were additionally considered.

**Figure 3.**
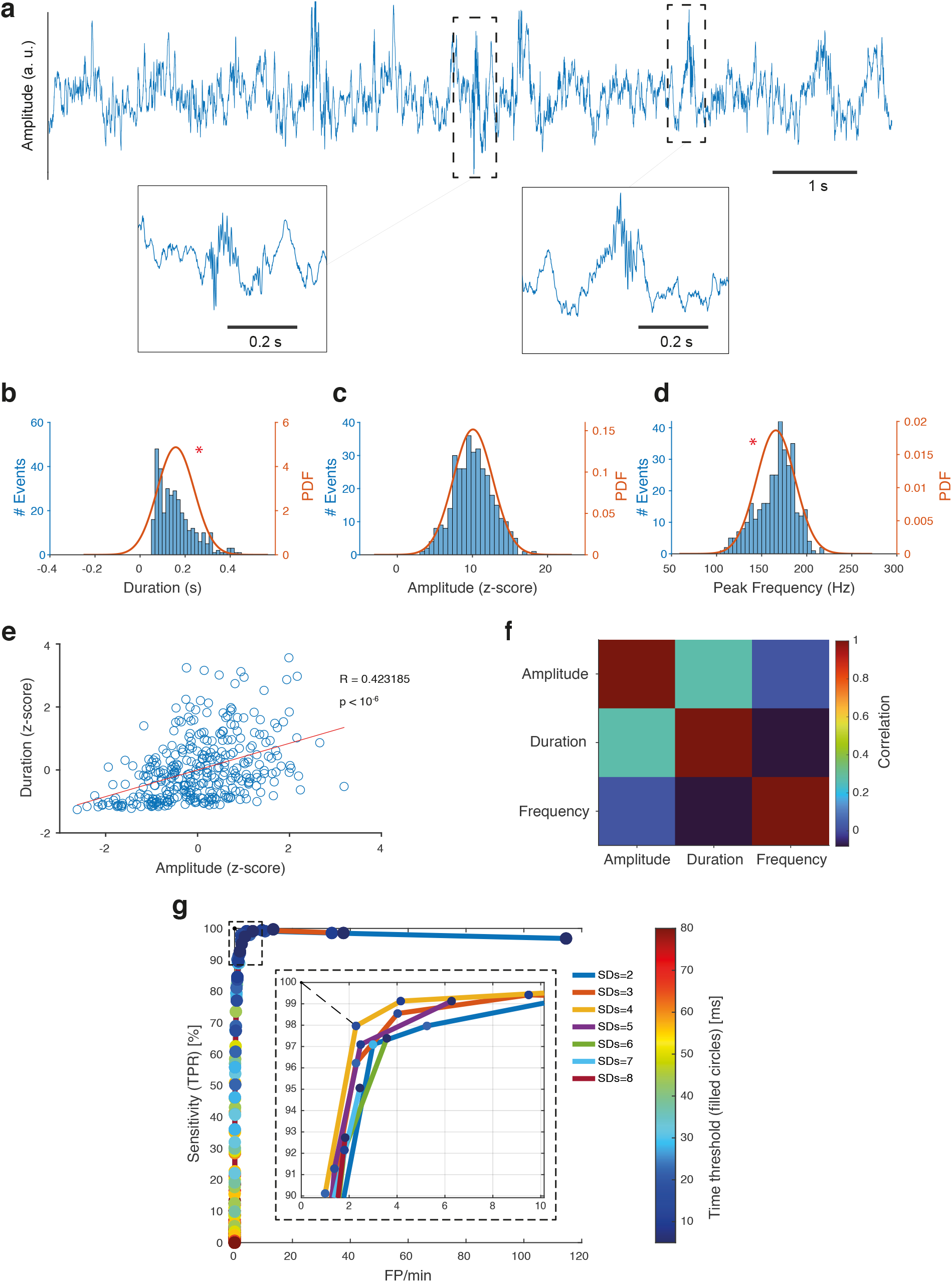
Characterization of real, pre-recorded ripple data (hc-3 dataset) and corresponding ROC curves for offline simulations of the plugin in Matlab. **a)** Example of an LFP segment. Two real ripple events are highlighted. **b)** Duration of ripple events (n=345 events, Kolmogorov-Smirnov normality test, null hypothesis rejected with p=0.0003; median=144 ms, interquartile range=111.2 ms). **c)** Amplitude (maximum value of the ripple RMS envelope) specified in SDs of the background noise (z-scores) (Kolmogorov-Smirnov normality test with p=0.9444; mean=10.07 z-units, SD=2.63 z-units). **d)** Peak frequency of ripple events (Kolmogorov-Smirnov normality test, null hypothesis rejected with p=0.0112; median=169.25 Hz; interquartile range=29.25 Hz). **e)** Scatter plot of amplitude *versus* duration of ripple events (Pearson’s corr. R=0.423185, Student’s t-test with p<10^−6^). The red line indicates a linear fit of the data. **f)** Visual Pearson’s correlation matrix for amplitude, duration, and frequency of ripple events. **g)** ROC curves for offline Matlab simulations of the plugin. The color bar and the respective color dots represent different time threshold values (ms). Various amplitude threshold values (specified in SDs) are represented by different curve line colors (blue: SDs=2, orange: SDs=3, yellow: SDs=4, purple: SDs=5, green: SDs=6, light blue: SDs=7, red: SDs=8). The dashed square is a zoom of the top left corner of the ROC graph. The dashed line indicates the closest point (97.9% TPR and 2.29 FP/minute) to the optimal reference.

### 2.5. Prior identification of candidate ripple events

The classification of ripple events in the hc-3 dataset was preceded by a prior identification of candidate ripple events. A modified version of a previously reported detection algorithm was applied (Taxidis et al., 2015). First, the LFP is bandpass filtered at 100-250 Hz using an FIR filter with a linear phase. The squared values of the filtered LFP are then computed, averaged with a zero-phase FIR filter (order 62), and standardized. Ripple events are initially detected when this standardized power average crosses the threshold of 1 SD (calculated over the entire data). Consecutive events separated by less than 50 ms are merged, and those with a final duration outside the 50-450 ms range are discarded. Finally, only the events with a peak power between 2 and 250 SDs are considered ripple candidates to be further approved or rejected by visual inspection.

### 2.6. Closed-loop setup to measure the round trip delay

To measure the round trip delay – the time from the ripple onset to the stimulation – we set up a closed-loop architecture constituted by an Intan RHD2132 headstage, the Open Ephys acquisition board (open-ephys.org), a BNC-to-HDMI conversion board, the Pulse Pal device (an open-source pulse generator, sanworks.io), and a computer (Intel i5-8300H, 2.3GHz, 12 GB RAM) running Windows 10 (**Fig. 4a**). The synthetic data was converted to a.flac audio file and played directly to an Intan headstage channel via custom-made cable and connectors. The acquisition board receives data from the headstage via SPI cable and streams it to the Open Ephys GUI through a USB 3.0 connection. A processing pipeline in the Open Ephys GUI was built to filter the incoming data in the 100-250 Hz frequency range before supplying it to the ripple detector plugin. Additional plugins were inserted into the processing chain to record (Record Node) and visualize (LFP Viewer) the data, as well as to communicate with the Pulse Pal hardware (Pulse Pal). The detection events fed the Pulse Pal module, triggering a brief digital pulse (single 5 V monophasic pulse, 100 μs duration) sent back to the Open Ephys acquisition board. The synthetic signal captured by the headstage, the moments of ripple detection, and the digital pulses generated by Pulse Pal were logged to compute the round trip delay.

**Figure 4.**
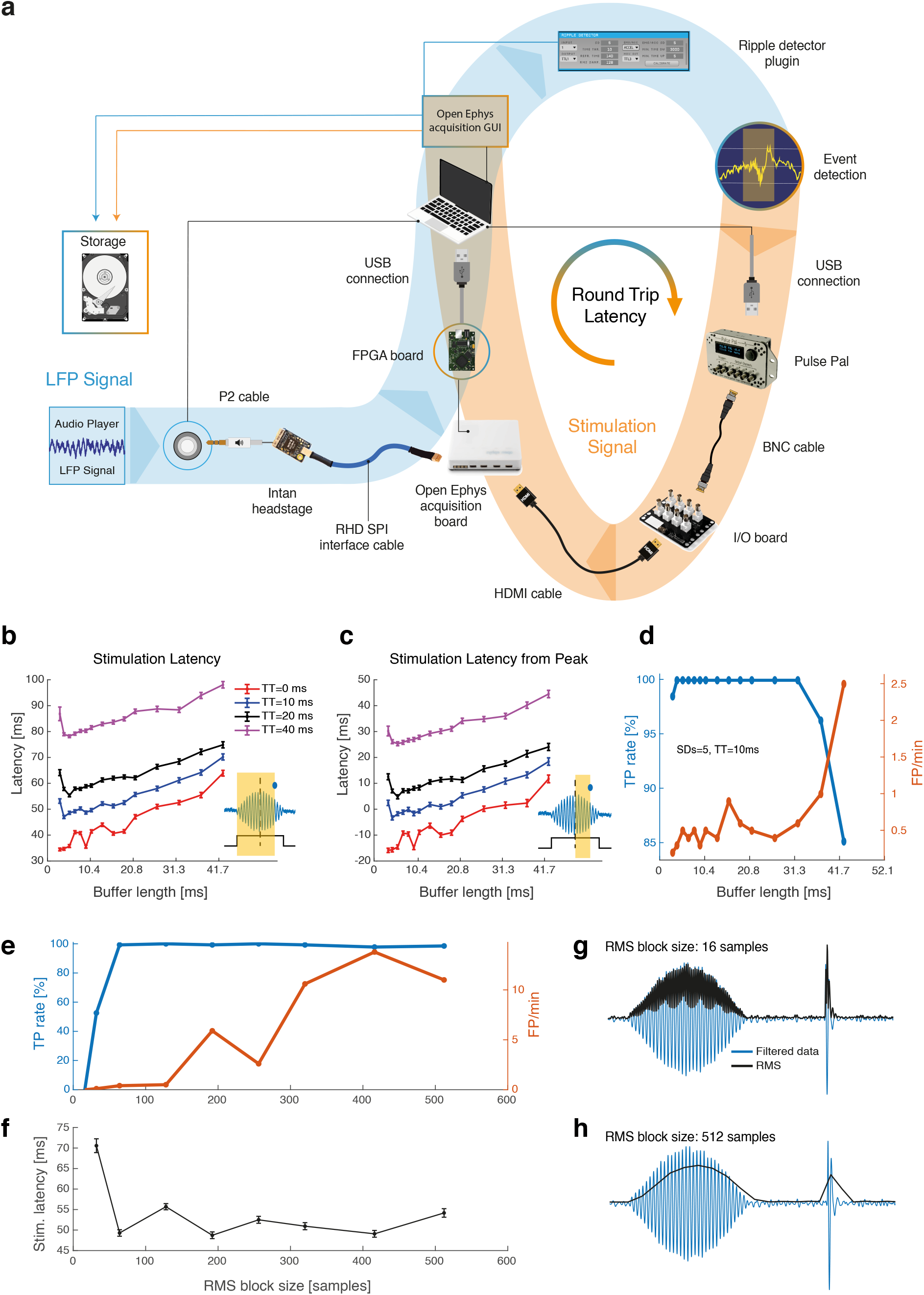
Performance evaluation of the closed-loop architecture for ripple detection and stimulation using synthetic data. **a)** Components of the closed-loop setup. Highlighted in blue is the LFP signal processing, and the stimulation signal processing is highlighted in orange. Synthetic LFP data containing simulated ripple events are converted to an audio file and streamed directly to the Intan headstage. The Open Ephys acquisition board sends the data to the computer running the Open Ephys GUI, where it is stored and filtered before it is forwarded to the developed ripple detector plugin. When a ripple is identified, the plugin produces an event that triggers a digital pulse via the Pulse Pal module. The acquisition board captured this stimulation pulse and stored it for further analysis (especially for latency computation). **b-c)** Round trip latencies of the closed-loop setup. Values are given regarding the ripple onset (**b**) and the ripple peak (**c**). Negative values in **c** indicate that the stimulation was applied before the ripple peak. Each line color represents different time threshold values (red=0 ms, blue=10 ms, black=20 ms, magenta=40 ms). We used the Windows Audio device with a sampling rate of 48 kHz. **d)** TPR and FPR for different buffer lengths. Amplitude and time threshold were fixed to 5 SDs and 10 ms, respectively. The blue line indicates the TPR (%), and the FPR (FP/minute) is indicated by the red line. **e)** Detection rates for different values of RMS block size. Blue line: TPR (%); red line: FPR (FP/minute). **f)** Round trip latency for different values of RMS block size. Filtered ripple and artifact data are illustrated with their respective envelopes for an RMS block size of 16 (**g**) and 512 samples (**h**).

### 2.7. Accelerometer data

The 3-axis accelerometer data used to validate the mechanism that silences ripple events after detecting movement was obtained from a public database available at the buzsakilab.nyumc.org/datasets/ website (Petersen, Hernandez and Buzsáki, 2020). We selected the TingleyD dataset (Tingley and Buzsáki, 2018), folder D12, session DT12_4032um_2232um_190806_090411. The accelerometer data segment was extracted from the auxiliary.dat file.

### 2.8. TPR and FPR

Three indices were calculated to characterize the detection performance: TPR, FPR, and round trip latency. For both the synthetic and pre-recorded real data, the TPR for a particular set of parameters (e.g., amplitude and time thresholds, RMS block size, buffer length) is given by the fraction between the sum of ripple events correctly detected and the total number of events previously classified as ripples (equation **1**). In particular, a detection event is considered in the TPR computation when it occurs during the time window that defines the ripple duration. For the synthetic data, the duration of each ripple is intrinsically known since the simulation of the corresponding event was produced under previously established conditions. Regarding the real data, the time window that constitutes the boundaries of a ripple activity was defined during the prior classification of candidate events. In other words, for un-merged ripples, the duration corresponds to the period in which the power average is above the 1 SD threshold (session **2.5**). For merged events, the duration comprises the onset of the earlier event and the end of the latter. Two or more consecutive detections of the same ripple event were considered once for the TPR calculation. The FPR was computed as the sum of the detection events in a non-ripple period divided by the total recording duration in minutes (equation **2**). We opted to provide the FPR as the number of false detections per unit of time because the determination of a denominator that represents the total number of false events (false ground events), albeit it could be defined according to some criteria, may generate a tricky, barely tangible result when the outputs of different experiments are compared.

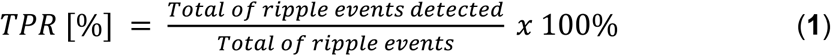

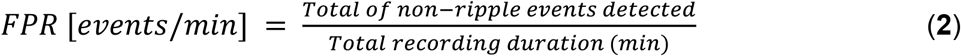

## 3. RESULTS

The ripple detector module contains a straightforward, user-friendly graphical interface (**Fig. 1a**) that allows investigators to select the input and output channels, adjust the detection thresholds and set a few more detection parameters, such as the number of samples for RMS calculation and the refractory time (period after detection during which new events are blocked). The refractory time is particularly important to avoid redundant output events caused by continued ripples. By clicking the “Calibrate” button in the graphical user’s interface (GUI), investigators can redefine the detection threshold by recalculating the baseline mean and standard deviation. Additionally, the embedded movement detector prevents ripple events from being raised when the animal is most active, reducing inappropriate stimulation due to movement artifacts. Because the ripple detector module is open-source, users with sufficient programming skills in C++ can modify the original source code to add new features and address specific requirements of their research.

### 3.1. Offline simulation in Matlab with synthetic data

To evaluate the performance of the ripple detector, the detection algorithm was first implemented in Matlab and tested with synthetic ripple data. Synthetic data was generated by injecting three types of simulated events in a background pink noise process: ripples, fast ripples, and spikes. Initially, to validate the effects of changing the thresholds in the detection performance, the amplitude threshold was fixed at 5 standard deviations (SDs) above the mean, and we varied the time threshold from 0 to 80 ms. As expected, higher time threshold values reduce the false-positive rate (FPR) by avoiding the detection of spike events but also decrease the algorithm’s sensitivity to detect shorter ripple events (**Fig. 2g**). Likewise, fixing the time threshold at 10 ms and increasing the amplitude threshold produced a similar result (**Fig. 2h**). Increments in the amplitude threshold resulted in the correct rejection of the residual power of filtered fast ripples, but they also limited true detections by hampering RMS crossing and time threshold attainment. Lastly, multiple simulations with different pairs of threshold values were conducted, and some of the detection rates obtained were strictly close to the reference point of 100% true positive rate (TPR) and 0% FPR in the receiver operating characteristic (ROC) graph (**Fig. 2i**).

Amplitude (maximum value of the ripple RMS envelope) of simulated ripple events specified in SDs of the background noise (Kolmogorov-Smirnov normality test with p=0.7856; mean=20.11 z-units, SD=1.07 z-unit). **d)** Peak frequency of ripple simulated events (Kolmogorov-Smirnov normality test with p=0.93; mean=201.71 Hz; SD=23.81 Hz). **e)** Scatter plot of amplitude *versus* duration of simulated ripple events. The red line indicates a linear fit of the data. A small correlation was observed (Pearson’s corr. R=0.236423, Student’s t-test with p=0.005078). **f)** Visual Pearson’s correlation matrix for amplitude, duration, and frequency of simulated ripple events. **g)** ROC curve for different time threshold values (amplitude threshold fixed to 5 SDs). The color bar and the respective colored dots represent the time threshold values (ms). The x-axis is the same as in graph **h** (FP/minute). **h)** ROC curve for different values of amplitude threshold (time threshold, TT, fixed to 10 ms). The color bar and the respective color dots represent the amplitude threshold values (specified in SDs). **i)** ROC curves for multiple simulations with different pairs of threshold values (amplitude and time thresholds). The color bar and the respective color dots represent the time threshold (ms). Different amplitude thresholds (specified in SDs) are represented by different curve line colors (blue: SDs=2, orange: SDs=3, yellow: SDs=4, purple: SDs=5, green: SDs=6, light blue: SDs=7, red: SDs=8). The dashed square is a zoom of the top left corner of the ROC graph (the optimal cut-off point), where the TPR is maximum (100%), and the FPR is minimum (0 FP/minute). The closest point achieved corresponds to 100% of TPR and0.1 FP/min (time threshold of 6 ms and amplitude threshold of 6 SDs).

### 3.2. Offline simulation in Matlab with real data

Different ripple events share common oscillatory features, but they are heterogeneous in duration, power, shape, frequency range, and frequency distribution (Nguyen et al., 2009; Fernández-Ruiz et al., 2019). Although synthetic data provide a reasonable idea about the plugin’s behavior and performance, it is essential to validate the detection algorithm over real ripple recordings. We downloaded a dataset publicly available on the CRCNS website that contains real hippocampus recordings from a Long-Evans rat to test the plugin’s detection (hc-3 dataset). Local field potential (LFP) data from the superficial layer (stratum oriens) of the CA1 hippocampal region was selected from a period when the animal was asleep. To label the ripple events for TPR calculation, we first ran a prior detection algorithm based on previous work (Taxidis et al., 2015). After this initial ripple identification, experts visually inspected and curated each candidate event to guarantee that no spurious oscillation (e.g., movement or chewing artifacts) would be classified as a true ripple event. The end-up number of confirmed ripples was 345 (11 events per minute). Then, multiple simulation sessions of the developed algorithm were conducted in Matlab with different amplitude and time thresholds to obtain the corresponding ROC curves (**Fig. 3g**). Many combinations of threshold values resulted in satisfactory detection rates, and the closest to the optimal cut-off point of the ROC graph, considering the Euclidean distance of the normalized coordinates, was 97.9% TPR and 2.29 FP/min (amplitude threshold of 4 SDs and time threshold of 12 ms).

### 3.3. Real-time, closed-loop experiments using Open Ephys with synthetic data

Offline simulation offers an acceptable output of the detection algorithm when its purpose is to perform a preliminary validation of the theoretical concepts behind the implemented logic (e.g., adjusting sensitivity by changing the detection thresholds). The Matlab implementation of the detection algorithm does not mimic with fidelity some aspects of the recording setup with Open Ephys, such as small random fluctuations in the data buffer delivery, communication delays between the hardware components, and contamination by different sources of electrical noise. Therefore, some performance indicators are more reliable when obtained from an experimental scenario similar to that used during real experiments. One of these performance indicators is the round trip latency: the delay between the ripple onset and the stimulation/manipulation command in closed-loop systems. In most experiments that sought to interfere in neuronal circuits in a real-time, ripple-dependent manner, the round trip delay must be as minimum as possible without impacting negatively and significantly the detection rates (TPR and FPR). Those experiments commonly include SWR disruption itself (Girardeau et al., 2009; Jadhav et al., 2012 and many other studies shown in Table I), but also the manipulation of different brain areas associated with ripple activity and its biological functions (Maingret et al., 2016).

A closed-loop architecture was assembled to measure the round trip latency (**Fig. 4a**), and the LFP of the living animal was replaced by the synthetic data converted to an audio signal. The data acquisition system recorded the stimulation command back. Because the time threshold theoretically imposes an intrinsic algorithmic delay to the ripple detection, we initially sought to confirm the impact of increasing this parameter on the round trip latency. In this case, the amplitude threshold was fixed to 5 standard deviations above the mean. Increments in the time threshold are followed by an almost equivalent increase in the latency (**Fig. 4b**,**c**). Buffer length also plays an important role in the round trip delay. This parameter can be manually adjusted in the Open Ephys GUI at runtime, and it was observed that higher buffer lengths increased the round trip latency, probably due to the intrinsic additional time to accumulate more data samples. By setting the time threshold to 0 ms, a mean delay of around 35 ms (34.5 ± 0.5 ms) from the beginning of the ripples (**Fig. 4b**) or around 15 ms (15.9 ± 0.8 ms) before the ripple peak (**Fig. 4c**) was obtained when buffer length was minimal (3 ms). It is worth noting that, for the synthetic data, the ripple power peak matches half the duration of the event. Buffer length also affects on the detection rates, worsening TPR and FPR as it approaches the maximum value of 42.7 ms (**Fig. 4d**).

The detection algorithm operates with the ripple-band filtered signal envelope given by the RMS values of sequential blocks of samples. To verify how the quality of the RMS contour may affect latency and detection rates, the plugin’s performance was checked for different values of RMS window lengths (number of samples used to calculate this metric). The amplitude threshold was then fixed to 5 SDs, the time threshold to 10 ms, and the buffer length to 21.3 ms. Shorter RMS sample blocks (i.e., 16 or 32 samples) produce a fast-tracking envelope that constantly resets the algorithm before the time threshold is fully accomplished, resulting in lower TPR (**Fig. 4e, g**). On the other hand, extremely large RMS blocks (i.e., 512 samples) extend spike duration and contribute to increasing FPR (**Fig. 4e, h**). Regarding the latency, we initially thought that larger RMS blocks would result in higher detection (and thus stimulation) delay. However, the results do not indicate a rising latency trend following increments in the RMS length (**Fig. 4f**). Instead, smaller blocks seem to impose an additional delay due to the resetting mechanism.

It is worth noting that the RMS block length must be adjusted according to the sampling frequency to avoid the drawbacks of selecting extreme values. We suggest 2.5 RMS values per period of the lowest frequency in the ripple band for an acceptable envelope. Because 100 Hz was the lowest ripple-band frequency used in this study (period of 10 ms), it corresponds to 4 ms of data comprised in each RMS value. With a sampling frequency of 30 kHz (the same used to build the synthetic data), the recommended RMS length would include 120 samples. For other values of sampling frequency (Fs) and lowest ripple-band frequency (Lf), the suggested RMS block size can be calculated as:

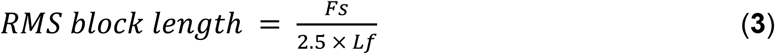

Based on the results obtained with the synthetic and real datasets tested in this work, our recommendation of parameters for general ripple detection is 5 SDs for amplitude threshold, 10 ms for time threshold, 21.3 ms for the buffer length, 140 ms for the refractory time, and an RMS block length that follows equation (**3**). Those values led to 97% of TPR and 2.48 FP/min for the real data. Regarding the synthetic data, the same detection parameters resulted in 100% of TPR, 0.5 FP/min, and a mean round trip latency of 55.7 ms from the ripple onset or 5.3 ms from the ripple peak. Although it is possible to decrease the latency even more by reducing the amplitude and time thresholds, investigators must be aware of the trade-offs concerning the detection rates, especially the FPR. Finally, we recommend executing prior recordings to optimally adjust the detection parameters according to the experimental requirements and conditions. The File Reader plugin from the Open Ephys GUI can be used for the initial calibration of the ripple detector parameters. For this purpose, Matlab scripts that convert the recorded data to a File Reader compatible file are available at github.com/open-ephys/analysis-tools.

### 3.4. Disabling ripple detection by monitoring movement in real-time

In vivo experiments involving long-lasting electrophysiological recordings and real-time, ripple-triggered manipulations may benefit from an automatic mechanism that prevents events from being raised when movement is detected. Especially when stimulation must be performed only during sleep or immobile wakefulness, the ripple-band power augmentation caused by movement artifacts may generate spurious events that interfere with the experimental protocol. To validate the blockage of ripple events based on movement detection, we used real 3-axis accelerometer data publicly available (Petersen, Hernandez and Buzsáki, 2020). The accelerometer data was converted to an appropriate file and loaded in Open Ephys GUI with the File Reader module. The detection algorithm first calculates the magnitude of the acceleration vector based on its 3-axis filtered data (100-250 Hz) and then proceeds with RMS computation, calibration, verification of threshold crossing, and, if necessary, event blockage (detailed description in section **2.2**). As expected, higher amplitude thresholds decrease the plugin’s ability to detect small changes in acceleration (**Fig. 5b** compared to **Fig. 5e**). This decreased sensitivity favors experimental scenarios in which stimulation should be applied despite small movements during brief periods of micro-arousals (Lima et al., 2019), for example. It was also observed that an increase in the time threshold permits only longer, persistent speed-changing movements that disable ripple events (**Fig. 5b** compared to **Fig. 5f**). Finally, increments in the minimum period of immobility required to restore the plugin’s capacity to trigger stimulation result in longer-lasting blockage (**Fig. 5a, b, c, d**). The minimum period of immobility controls the extension of the silencing, and it can be adjusted, if necessary, to prevent LFP transients during short periods of immobile wakefulness from triggering stimulation.

**Figure 5.**
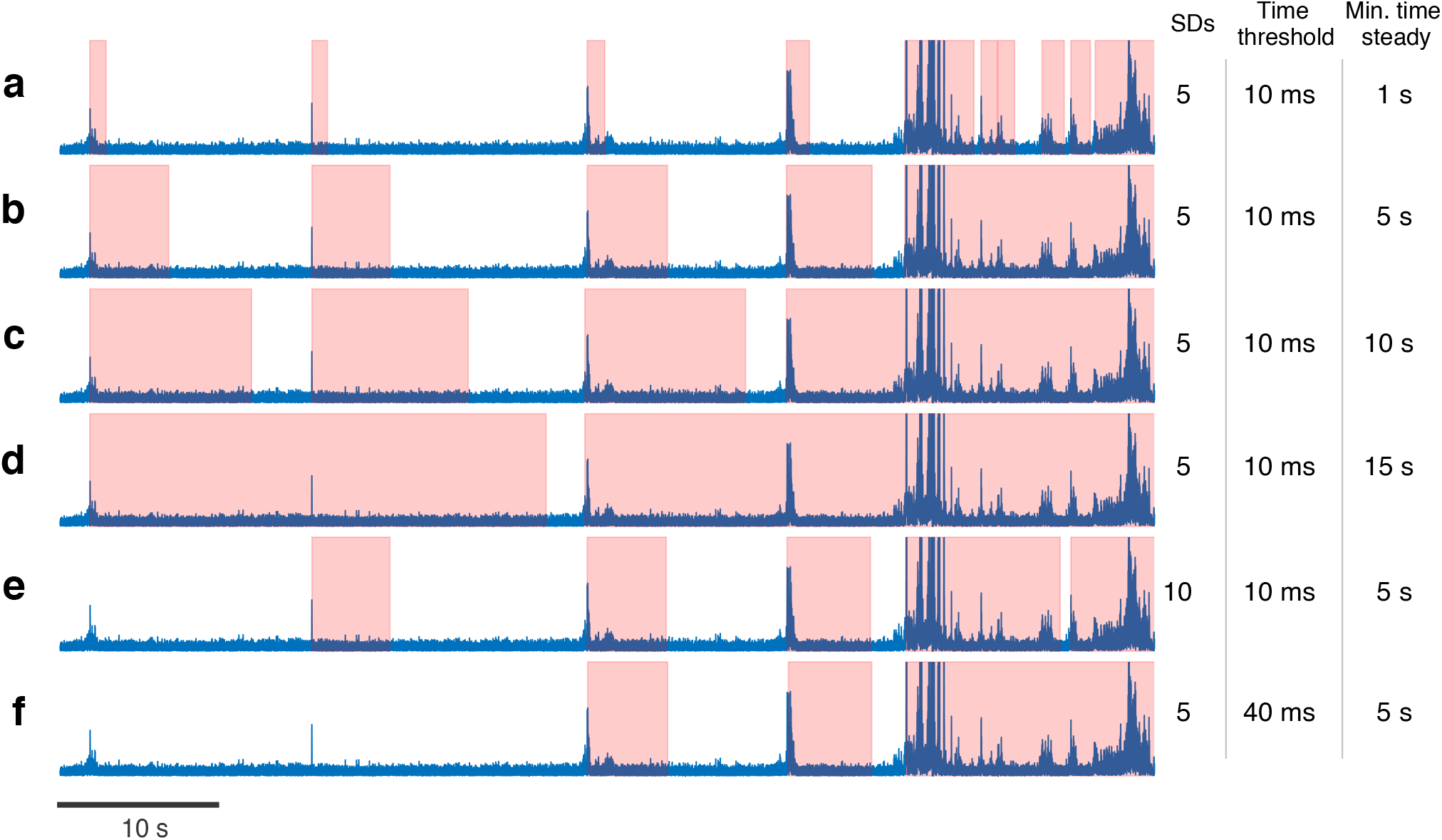
Movement detection disables ripple events. **a-f)** Six scenarios of event blockage produced by different combinations of time threshold, amplitude SDs, and minimum immobility period. Blue lines: magnitude of the accelerometer vector. Areas highlighted in red: periods in which ripple events are silenced after movement is detected.

### 3.5. Multichannel ripple detection

Electrophysiological recordings of the hippocampus are commonly carried out using multiple electrode arrays. Ripples are commonly observed in the CA1 pyramidal layer, and their shape varies with the depth of the implant (Buzsáki, 2015; Ramirez-Villegas, Logothetis and Besserve, 2015). In this scenario, a multichannel ripple detection logic offers the possibility to enhance the detector’s accuracy. We verified the combined ripple detection performance for amplitude threshold values using three simultaneous CA1 recordings from hc-3, the same dataset used in section **3.2**. The corresponding electrode tips were vertically 40 μm apart from one another, and the silicon probe shank was aligned parallel to the septo-temporal axis (45º parasagittal) (Mizuseki et al., 2014). Because the audio output is limited to 2 channels, the File Reader module was used to stream the 3-channel data. An Open Ephys processing chain was set up to combine the outputs of 3 ripple detector modules in a boolean AND logic, each detector applied over a different single channel (**Fig. 6a**). The Logic Gate module, a third-party plugin developed for the Open Ephys GUI, was used for the logical processing of the individual outputs (Buccino et al., 2018). As expected by using the AND expression, the TPR of the combined strategy was limited by the lowest sensitivity of an individual detector (**Fig. 6b**). Moreover, by triggering an event only when all the individual detectors identify a ripple simultaneously, the global FPR can be reduced (**Fig. 6c**). We intend to implement a customizable, multichannel algorithm internally in future releases of our ripple detector.

**Figure 6.**
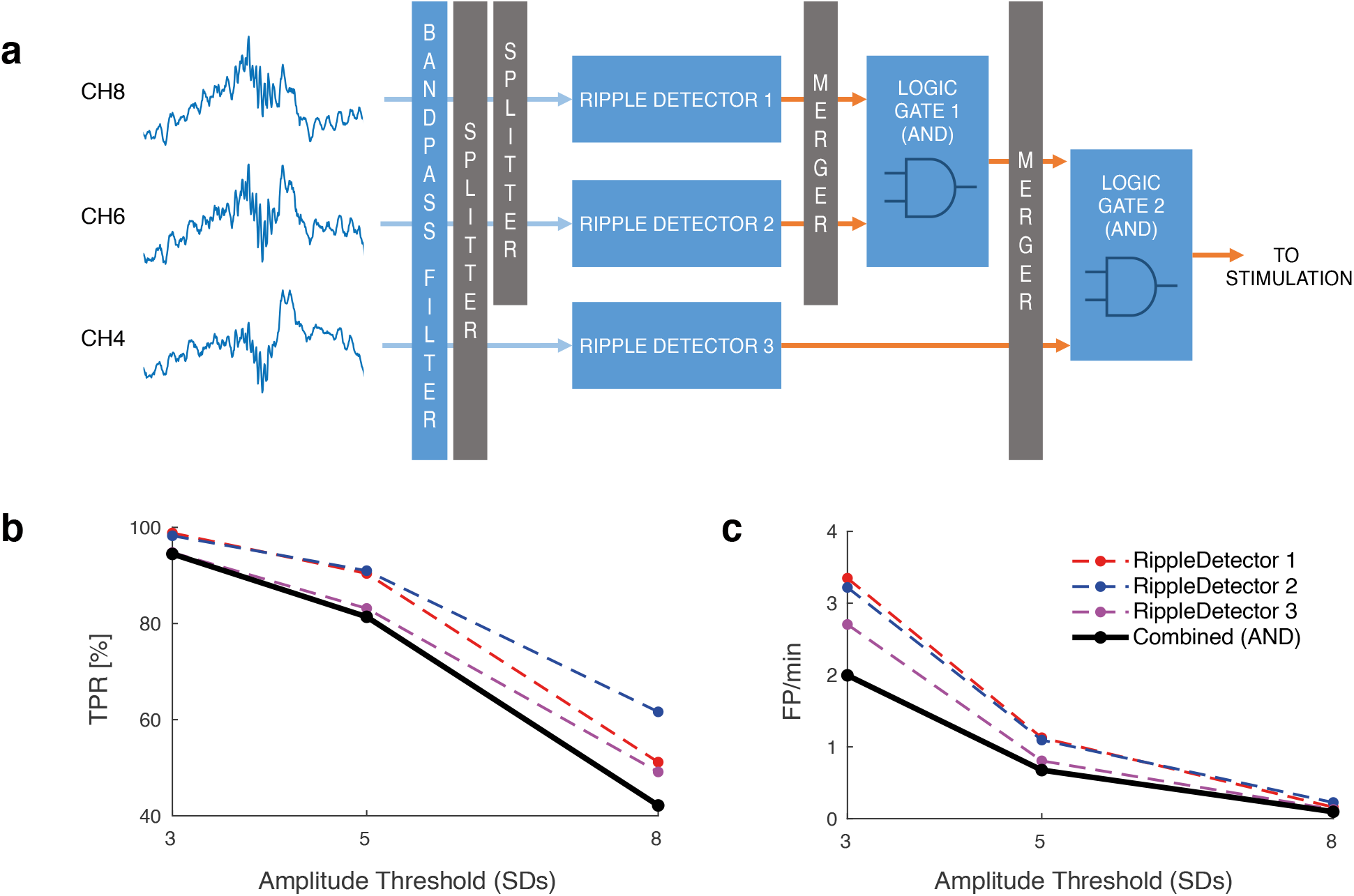
Multichannel ripple detection analysis. Three simultaneous CA1 recordings from the hc-3 dataset were used (a distance of 40 μm between the recording sites). **a)** Schematic representation of the multichannel ripple detection processing chain built in the Open Ephys GUI. **b)** TPR for single and multichannel detections. Different amplitude thresholds were tested (3, 5, and 8 SDs). Each single-channel detection is represented by one dashed line (red=ripple detector 1, channel 8; blue=ripple detector 2, channel 6; magenta=ripple detector 3, channel 4). The continuous black line shows the combination of the 3 detector outputs using a boolean AND logic (ripple detector 1 AND ripple detector 2 AND ripple detector 3). **c)** FPR for single and multichannel detections.

## 4. DISCUSSION

We presented the development and the validation of a ripple detector plugin integrated with the Open Ephys data acquisition platform. The plugin has a user-friendly graphical interface that allows investigators to adjust some detection parameters to balance round trip delay versus detection accuracy. Furthermore, its open-source nature allows community-aided improvements, and the adaptation of the internal algorithms for particular needs. In addition, to reduce spurious stimulation and enhance the quality of the experiments that investigate the importance of SWRs to memory and other processes, we developed a built-in, automatic on/off mechanism based on movement detection.

The results suggest an important interaction between the detection parameters and the performance trade-offs (i.e., round trip delay and detection accuracy). An increase in the time threshold, for instance, contributes to reducing the false-positive rate while imposing more delay to generate the stimulation. The consequences of varying other parameters than those directly related to the detection thresholds (i.e., buffer length and RMS block size) must also be considered. As shown in this work, the RMS block size affects the quality of the envelope and should be adjusted to avoid both the signal shifting and the over-representation of ripple cycles. Likewise, shorter buffer lengths may not contain the appropriate number of samples to calculate the RMS, while the longer ones are associated with increased stimulation delay. The mutual interaction between the parameters and the detection trade-offs are probably unknown by the investigators, as suggested by the absence of performance reports (Table I), and can impair their experimental protocols and the associated results. In a worst-case scenario, the effects observed in experimental groups may result from the excessive intervention itself due to a detection algorithm with low specificity. Similarly, rapidly-triggered but insufficient manipulations due to a low-sensitivity detection may contribute to the absence of effect between groups. It is worth noting that the choice between accuracy and latency is intimately connected to the hypothesis being tested and the experimental design. Depending on the study, it may be acceptable or even desirable to prioritize one of those features over the other without major impairments. In any case, the detection system must be previously well-characterized to the setup being used.

Another important aspect we addressed was the need to avoid movement artifacts from triggering the stimulation/manipulation system. The integrated algorithm that prevents ripple events from being raised during mobility periods reduces false positives and the effects of inappropriate stimulation. Furthermore, because it is an automatic mechanism, it exempts investigators from the continuous inspection and manual on/off intervention, supporting the execution of simultaneous recordings in different animals.

One advantage of implementing movement detection based on the accelerometer or EMG data, as proposed in this work, is that those signals are usually already present in memory and sleep studies to determine the animal states (wakefulness or sleep) (Boyce et al., 2016; Koike et al., 2017; Lima et al., 2019). However, different strategies have been used throughout the literature to prevent movement artifacts from resulting in false detection. One of those strategies is to implement a speed filter based on the live image of the behaving animal to block the generation of stimulation events when the animal’s speed crosses a predefined threshold (around 5 cm/sec, Table I). The limitations of this approach reside in the inability to detect movement artifacts while the animal’s position remains unchangeable. That includes chewing, grooming, and headstage knocking against the box, for example. Another strategy is based on positioning a reference or control electrode outside the hippocampal formation to detect common LFP transients that leak the ripple frequency range (Talakoub et al., 2016; Oliva et al., 2020; van de Ven et al., 2016). The simultaneous occurrence of these high-frequency transients in a non-hippocampal region is used to cancel the “ripple” detection in the hippocampal recording channel. Besides demanding an additional implant, this scheme may present a disadvantage when used in experiments of selective manipulation during sleep. Since the LFP of the control electrode may remain smooth during wake periods of low mobility, awake ripples are detected without distinction. With none or a few extra programming lines in the plugin’s source code, it is possible to use the control channel to turn off stimulation as it provides EMG data. Indeed, what the movement detection algorithm generically does is to verify significant changes in the RMS of the selected channel using the logic of threshold crossing already described. An alternative approach is to use an additional plugin instance on the control electrode channel to implement a cortical “NOT” and block ripple events.

The lowest round trip latency mean reported in our closed-loop experiments was 34.5 ± 0.5 ms for a buffer length of 3 ms, amplitude threshold of 5 SDs, and no time threshold (**Fig. 4b**, red line). This latency corresponds to 15.9 ± 0.8 ms before the ripple peak. Apart from the time threshold, that imposes an intrinsic algorithmic delay (**Fig. 4**), one of the most significant components of the round trip latency is the USB communication that links the computer executing the Open Ephys GUI to the data acquisition system and the Pulse Pal module. This USB overhead is around 20 ms (Siegle et al., 2017). Investigators interested in reducing the round trip latency can achieve it by lowering the amplitude threshold, the time threshold and/or the buffer length, but they should keep in mind the possible consequences on the detection rates. Because the plugin is open-source, it is also possible to integrate it with devices that accept communication protocols faster than USB (i.e., Ethernet or PCI express). For example, Neuropixels probes (which use a PCIe-based acquisition system; Jun et al., 2017; Putzeys et al., 2019) readily integrate with the Open Ephys GUI and would therefore be immediately compatible with our ripple detection system. Additionally, other initiatives (Open Ephys, 2016) and a new generation of the Open Ephys acquisition system are being developed to support submillisecond closed-loop systems with PCI express (open-ephys.github.io/onix-docs/). Although the USB overheads impose an intrinsic, considerable delay, the final latency is acceptable for many applications that are not usually accessible to neuroscientists (Siegle et al., 2017). That includes brain-machine interface systems and experiments involving ripple-triggered manipulations, whether in the hippocampus or other brain areas.

For future work, we intend to implement multichannel detection inside the plugin’s structure and evaluate how additional channels impact the round trip latency. In addition, testing different boolean equations is important to achieve better accuracy. In experiments that demand the operation of the plugin for long periods, the parameters calculated in the calibration stage might become less representative over time. Although the calibration button in the plugin’s GUI allows the recalculation of RMS mean and standard deviation, it would be useful to develop and test an algorithm for continuously estimating these parameters. Finally, since the developed plugin is based on RMS threshold crossing, we believe it can be adapted to detect other electrophysiological events, whether physiological (e.g., thalamo-cortical spindles) or pathological (e.g., interictal epileptiform discharges – Gelinas et al., 2016). While we have not conducted further analysis on it, it is possible to detect spikes or fast ripples in the synthetic data only by changing the cutoff frequencies of the bandpass filter (**Fig. S1**).

## ACKNOWLEDGMENTS

This work was supported by the National Council for Scientific and Technological Development (CNPq, Brazil - 422911/2021-6), the Minas Gerais Research Foundation, FAPEMIG (Universal APQ-00246-21), the São Paulo Research Foundation, FAPESP (2015/25275-8 and 2016/17882-4), and the International Society for Neurochemistry (CAEN-1B and CDG programs). We would like to thank Flávio A. G. Mourão for the discussions during the development of this work, Alysson R. da Silva for the initial support on programming and Josh Siegle for the valuable comments on this manuscript.

## COMPETING INTERESTS

The authors declare no competing interests.

## Notes

https://github.com/bmsousa91/RippleDetector

### Competing Interest Statement

The authors have declared no competing interest.

