## Supplemental Figure 1 for "An open-source, ready-to-use and validated ripple detector plugin for the Open Ephys GUI"

**SUPPLEMENTARY**

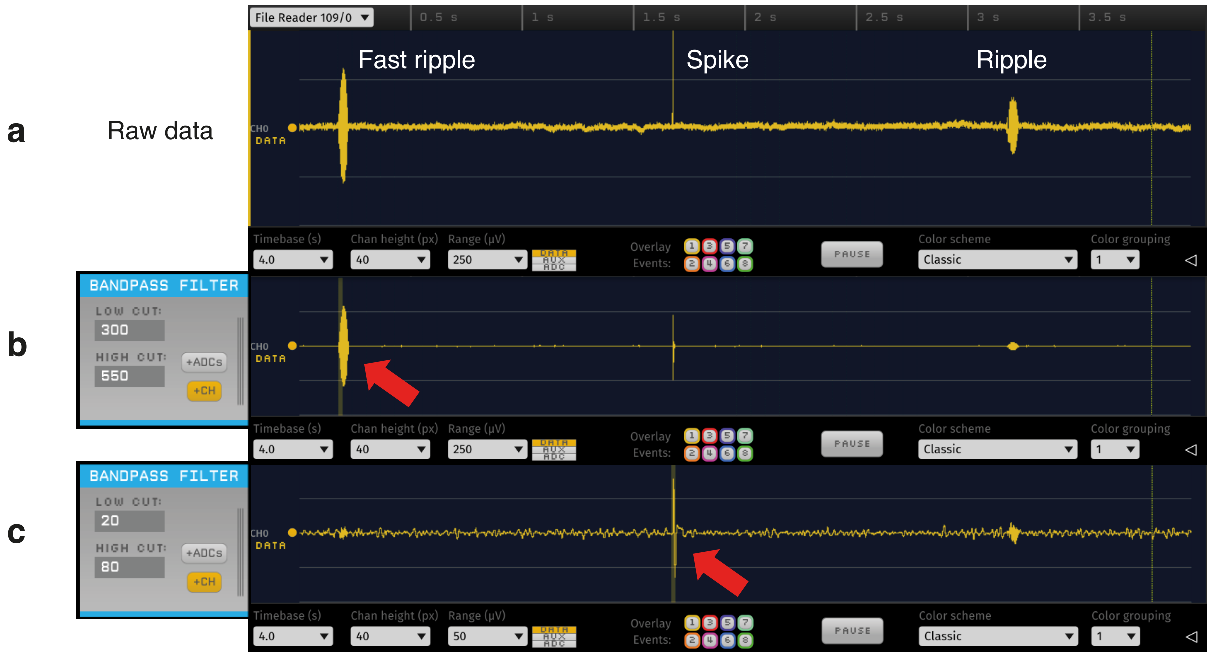


**Figure S1.** Example of spike and fast ripple detection in the Open Ephys GUI using the developed plugin. **a)** Segment of synthetic raw data. **b)** The same segment of synthetic data in **a**, but filtered in the 300-550 Hz frequency range to prioritize fast ripples. The red arrow indicates the detection of a fast ripple event. Detection parameters: amplitude threshold=5 SDs; time threshold=10 ms; RMS block size=128 samples; buffer size=1024 samples. Accuracy: 100% TPR (119 events – events during the calibration step were ignored) and 0 FP. **c)** The same segment of synthetic data in **a**, but filtered in the 20-80 Hz frequency range to prioritize spike events. The red arrow indicates the detection of a spike event. The detection parameters are the same as in **b**. Detection accuracy: 100% TPR (132 events – events during the calibration step were ignored) and 0 FP.
